# Current challenges for epitope-agnostic TCR interaction prediction and a new perspective derived from image classification

**DOI:** 10.1101/2019.12.18.880146

**Authors:** Pieter Moris, Joey De Pauw, Anna Postovskaya, Sofie Gielis, Nicolas De Neuter, Wout Bittremieux, Benson Ogunjimi, Kris Laukens, Pieter Meysman

## Abstract

The prediction of epitope recognition by T-cell receptors (TCRs) has seen many advancements in recent years, with several methods now available that can predict recognition for a specific set of epitopes. However, the generic case of evaluating all possible TCR-epitope pairs remains challenging, mainly due to the high diversity of the interacting sequences and the limited amount of currently available training data. In this work, we provide an overview of the current state of this unsolved problem. First, we examine appropriate validation strategies to accurately assess the generalization performance of generic TCR-epitope recognition models when applied to both known and novel epitopes. In addition, we present a novel feature representation approach which we call ImRex (interaction map recognition). This approach is based on the pairwise combination of physicochemical properties of the individual amino acids in the CDR3 and epitope sequences, which provides a convolutional neural network with the combined representation of both sequences. Lastly, we highlight various challenges that are particular to TCR-epitope data and that can adversely affect model performance. These include the issue of selecting negative data, the imbalanced epitope distribution of curated TCR-epitope datasets, and the potential exchangeability of TCR alpha and beta chains. Our results indicate that while extrapolation to novel epitopes remains a difficult challenge, ImRex makes this feasible for a subset of epitopes that are not too dissimilar from the training data. We show that appropriate feature engineering methods and rigorous benchmark standards are required to create and validate TCR-epitope predictive models.

## Introduction

For an adaptive immune response to be initiated, a T-cell receptor (TCR) on the surface of a T-cell has to recognize an immunogenic non-self or altered-self peptide (epitope) presented in the context of a major histocompatibility complex (MHC) molecule on the surface of another cell. This system has to be able to recognise a large variety of epitopes derived from rapidly evolving pathogens or malignant cells locked in an evolutionary arms race. The recognition of this diverse set of epitopes is made possible by the highly polymorphic MHC genes and the quasi-random somatic V(D)J recombination of the TCRs.

Understanding the patterns that govern successful TCR-peptide-MHC (pMHC) interaction and being able to predict the binding of any given TCR-epitope pair, would be an important step forward towards personalized healthcare. For example, knowledge on the specificity of the TCR repertoire could provide information on the progression of an infectious disease and aid in the assembly of effective vaccines, support the development of individually-tailored cancer immunotherapy and deliver new prospects in the research regarding autoimmune disorders.

Given the high intra- and inter-individual diversity of the TCR-pMHC interactions and the difficulty of mapping them through experimental means, computational methods offer a promising complementary approach to gain insight into the general patterns of interaction-driven activation.

Thus far, research has primarily focused on modeling MHC-epitope binding affinity and this has led to several reliable predictive tools (1–7). However, understanding the molecular underpinnings of TCR-epitope recognition has proven to be more challenging. This can be explained in part by the larger diversity of TCR sequences when compared to MHC alleles, especially within an individual. This diversity leads to a large, high-dimensional, and likely non-homogeneous search space, which necessitates sophisticated models that can efficiently extract the underlying distinguishing features from the highly variable molecular sequences. However, to train these models a large amount of high-quality data is required, which is currently still lacking because data generation can only happen at much smaller scales than for MHC data. Fortunately, more and more experimental data is constantly being generated and stored in public databases such as VDJdb, IEDB and McPAS-TCR (8–10).

In essence, these models need to be able to understand the processes that determine the affinity between a TCR and epitope on a molecular level. This process is rather complex, because even though sequence similarity seems to be a key factor, epitopes can be bound by varying TCRs (11–13), and conversely TCRs can also display cross-reactivity against multiple epitopes (14). Another complicating factor is that despite the advent of single cell TCR paired chain sequencing, the currently available TCR-epitope binding data still mostly consists of single-chain (usually beta-chain) data, while in reality both the alpha- and beta chains are thought to contribute to binding specificity. Indeed, recent studies have suggested that utilising both chains could increase the accuracy of predictive TCR-epitope models (15, 16).

Previous work has tackled the TCR-epitope recognition problem from an epitope-specific angle, and demonstrated that the amino acid sequences of the TCR’s CDR3 region contain relevant information to predict epitope recognition using epitope-specific models (11, 12, 15, 17, 18). These types of predictive models have now been made accessible to immunology researchers via web tools such as TCRex (19). One shortcoming of an epitope-specific approach is that a separate model needs to be trained for every epitope (or for a set of epitopes in the case of multi-class models (18)). Every model requires sufficient training examples of TCRs with the same epitope specificity, which is not always readily available, for example in the case of neoepitopes.

Generic sequence-based models that can predict the interaction between any TCR and epitope are the next frontier. However, recent studies have shown that while generic models can achieve a reasonable performance on interactions that involve epitopes that were already encountered during training, they cannot yet reliably extrapolate to novel, unknown epitopes (i.e. epitopes that are unseen during training) (16, 20). This is likely caused by the large heterogeneity in potential TCR and epitope sequences, and the limited amount of readily available training data that covers the sequence space. The problem of predictive models under-performing on new data due to a shift in the distribution of the inputs (i.e. a divergent set of TCRs and/or epitopes) is more generally referred to as covariate shift (21). It is crucial that appropriate methods are used to assess these different types of generalization performance, i.e. known-epitope and epitope-agnostic prediction, and that they are not conflated. These two approaches are inherently answering different questions: “*How well can we predict interactions involving known epitopes?*” versus “*How well can we predict recognition of novel epitopes?*”

Given the fact that detecting TCR-epitope recognition can be considered more complex than MHC-epitope recognition, at least in part due to the larger diversity in sequence pairing and the limited amount of available training data, training complexity constitutes even more of a limiting factor in this setting. This necessitates relevant feature encoding. Indeed, studies have shown that feature engineering is essential for the prediction of paratope-epitope binding (22) and that it can also improve the performance of models for more general bioinformatics problems (23). Moreover, when complex machine learning methods are applied to small datasets, the dangers of overfitting and memorization of examples pose an even bigger threat. In the case of TCR-epitope recognition, the potential of models to overfit on common epitope motifs should be assessed and reduced.

To tackle these issues, we present an approach to predict TCR-epitope recognition that was inspired by image classification tasks, where convolutional neural networks (CNNs) models have shown strong performance (24). Our method relies on a novel feature representation method that converts the molecular sequences into an interaction map, which combines the physicochemical properties of both interactors on the aminoacid level (Fig. 1). This preprocessing step avoids the creation of an internal embedding for each interacting molecule independently. We postulate that this feature representation provides a better representation of the binding context and in that manner can improve model performance.

**Fig. 1.**
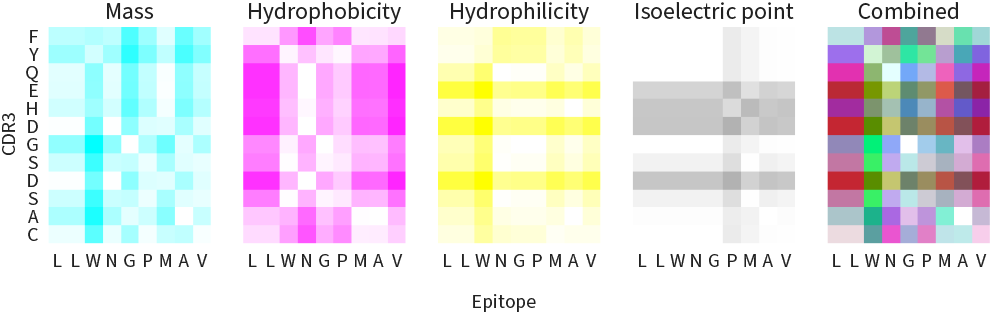
An interaction map of a CDR3 and epitope sequence encoded by four physicochemical properties using the absolute difference as the pairwise operator. The rightmost image is a combined representation of the four two-dimensional maps, using CMYK color encoding. As input to a CNN, the interaction map is supplied as a three-dimensional tensor.

Using such an interaction map as the input to a CNN differs from the prevailing approach seen in sequence-based neural network (NN), not only in the field of TCR-epitope prediction, but also for the modeling of other molecular interactions. Commonly, each sequence is supplied to the network as a distinct entity (i.e. in a separate input layer, before being combined later on in the network, or as a concatenated encoding (either at the sequence level or spread out over different channels of the input). (1, 3, 5–7, 16, 20, 25, 26). Consequently, the learning task not only consists of finding an internal representation that captures the properties of the amino acid residues in the two sequences individually, but also includes the combination of these higher-level features for the two interacting molecules in order to accurately predict their binding context.

In this paper, we train a CNN to predict the recognition of an epitope by a TCR, based on the amino acid sequences of the TCR’s CDR3 region and the epitope. First, the sequences are converted into an interaction map that represents a number of physicochemical properties of each amino acid pairing between the TCR and epitope. This interaction map is then fed into the CNN, which is tasked with learning how to extract the features that underlie the molecular binding process. We also contrast different strategies that can be used to tackle the the various biases that are particular to TCR-epitope recognition, such as negative data generation and epitope-bias.

## Methods

### Data

The data used in this study was collected from the August 2019 release of VDJdb (8). It consists of 75,474 curated pairs of TCR CDR3 alpha/beta sequences and their epitope targets, covering both MHC classes and three species (*Homo sapiens, Mus musculus* and *Macaca Mulatta*).

This dataset was reduced to 19,842 unique CDR3-epitope pairs by selecting only human TCR sequences (68,506), removing all spurious CDR3 sequences as defined by VDJdb (66,597), retaining only MHC class I entries (64,386), omitting all entries originating from the 10x Genomics demonstration study (27) (24,513), limiting the length of the CDR3 and epitope sequences to lie between 10-20 and 8-11 amino acids respectively (24,294), and finally removing any duplicate sequence pair. This *mixed chain* dataset was further split into an *alpha chain* (5,654 CDR3 alpha (TRA) sequences) and a *beta chain* (14,188 CDR3 beta (TRB) sequences) dataset. In some experiments the beta chain dataset was downsampled to address the epitope imbalance (see results for exact numbers). The mixed chain dataset contains 120 unique epitopes (60 and 118 for the alpha and beta chain datasets respectively), although their abundances are skewed; the majority of epitopes occur only in a few entries, whereas a small number of them appear in hundreds or thousands. An overview of the distribution of these sequences is given in Fig. 2.

**Fig. 2.**
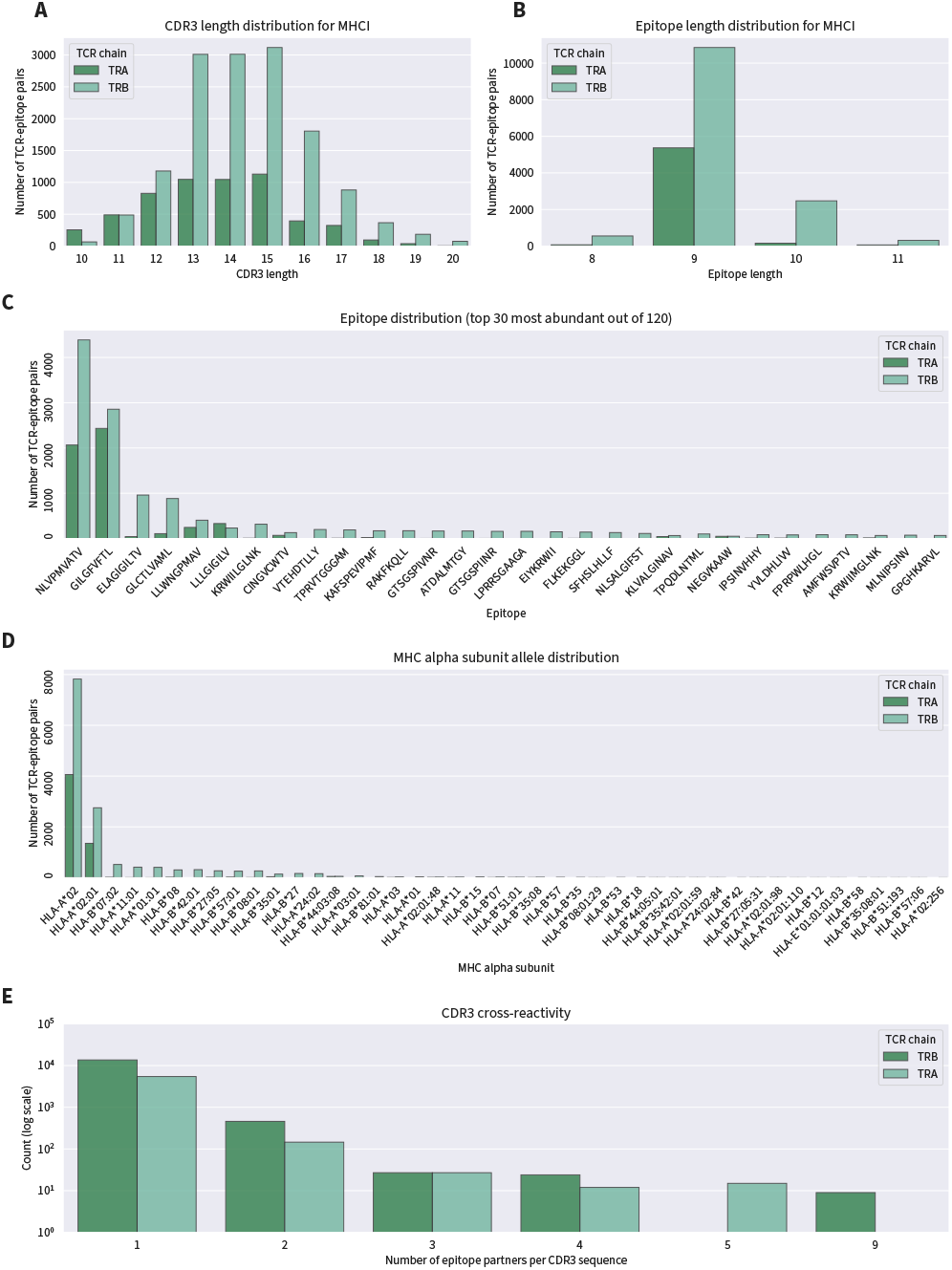
Overview of contents of the filtered VDJdb (mixed chain) dataset. A. Length distribution of TCR CDR3 sequence occurrences per TCR chain. B. Length distribution of epitope sequence occurrences per TCR chain. C. Top 30 most frequent epitopes.

Only MHC class I-derived pairs were used, since these made up over 90% of the dataset. The sequence length constraints were chosen in order to limit the amount of zero-padding required for our feature encoding and removed less than 1% of all pairs. We chose to exclude the 10x Genomics entries as the post-processing cut-offs for true epitope-TCR pairs is still a point of contention. Since our model only requires the CDR3 and epitope sequence as its inputs, all other metadata that is available in VDJdb was removed (e.g. V/J segments/genes). Lastly, we did not perform any filtering on the VDJdb confidence score.

An independent repertoire dataset consisting of 250,000 TRB CDR3 sequences, sampled from healthy TCR repertoires collected by Dean et al. (28), was used as reference data to generate negative examples in some experiments. The same size constraints as before were used and duplicate sequences were once again removed, resulting in 248,895 unique CDR3 sequences. This negative reference repertoire dataset showed minimal overlap with our filtered VDJdb dataset: only 749 sequences occurred in both.

The McPAS-TCR dataset (10) was used as an external test dataset (database release June 24, 2020). Only human MHC-I CDR3 sequences that were matched to an epitope via tetramer assays or peptide stimulation experiments were selected and the same length restrictions and deduplication as before were applied. In addition, any sequence pair present in the VDJdb dataset was removed. Afterwards, two subsets were created: the shared epitope subset containing 4,101 unique CDR3-epitope pairs covering a total of 47 epitopes, which were also present in the training data, and the unique epitope sub-set containing 736 pairs covering 11 unique epitopes, of which none were present in the training data.

### TCR-epitope feature encoding via interaction maps

We have designed a novel feature representation for sequencebased input data that summarises the binding context of the two interacting partners on the molecular level in terms oftheir physicochemcial properties. The model no longer has to learn an embedding for the individual interactors, which could help steer the model to focus on the actual interaction problem directly.

This encoding method can be thought of as the creation of an image or pixel map, and results in a 3-dimensional tensor that we will refer to as an interaction map. By lining up the sequence of the TCR and the epitope on the axes of a twodimensional matrix, every position within it corresponds to the pairwise combination (absolute difference) of a specific physicochemical property of the amino acids at that index of the two sequences. The map can have several channels, each corresponding to a different property, analogous to the colour channels of an RGB image. An example of an interaction map of a CDR3 and epitope sequence for four physicochemical properties is given in Fig. 1.

In our experiments, the interaction map for each CDR3-epitope pair was constructed as follows. First, each sequence in the pair was converted into a vector of physicochemical property values of each amino acid. Next, a matrix was computed that contained the pairwise absolute differences between the elements of the two vectors. One such twodimensional matrix was created for each of the following physicochemical properties: hydrophobicity, hydrophilicity, mass and isoelectric point (29). Every element in the matrix was then scaled between zero and one, based on the minimum and maximum possible values of each property. Next, the matrices were zero-padded on both sides to a dimension of 20 × 11 (*CDR3* × *epitope*). In the end, the matrices were combined into a three-dimensional tensor, where the length and width correspond to the two interacting sequences, and the depth (channels) to the physicochemical properties.

### ImRex CNN architecture

The architecture of the ImRex CNN was empirically determined and is depicted in Fig. 3 and follows the general design of CNNs for image classification. The input layer accepts interaction maps of dimensionality 20 × 11 × 4, where the first and second dimensions correspond to the amino acids of the TCR CDR3 and epitope sequences respectively, and the third (depth) corresponds to the different physicochemical properties. Next, there is the feature extraction part, which is made up of two identical units, each containing a series of 2D convolutional layers followed by a 2D max pooling layer and spatial dropout (25%). One is positioned after the first convolutional layer of each unit and the other after every dropout layer. The features extracted by the convolutional layers are then flattened and fed into a fully connected layer (32 neurons), followed by a single neuron output layer. All layers (except for the output layer) use an L2 regularization penalty of 0.01.

**Fig. 3.**
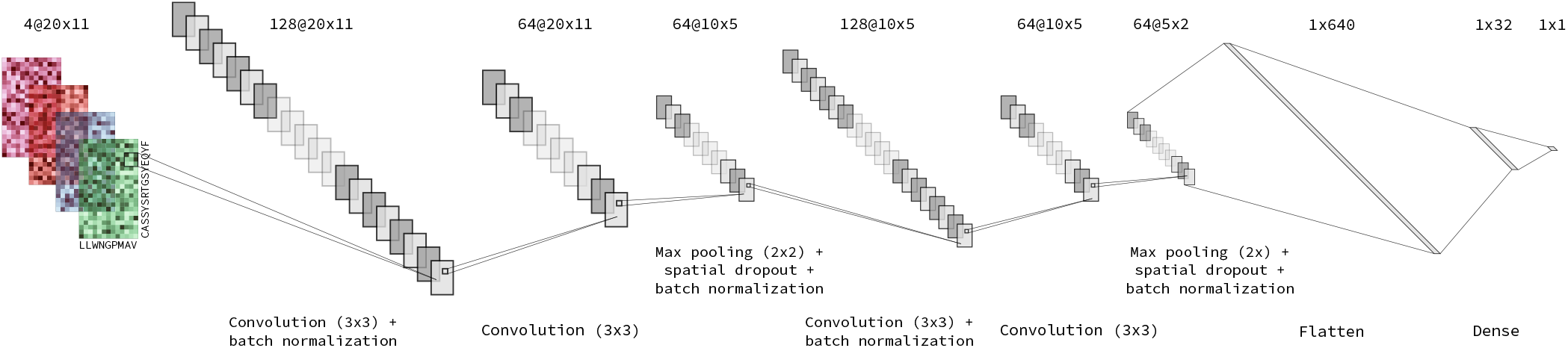
Architecture of the ImRex CNN. The four colours in the input layer represent the four different physicochemical properties used in the creation of the interaction map (hydrophobicity, hydrophilicity, mass and isoelectric point). Thin black lines originating from a smaller rectangle represent 2D convolutions, whereas other lines represent concatenation steps.

All neurons use rectified linear unit (ReLU) activation functions, except for the output neuron which relies on the sigmoid activation function. The convolutional layers use 3 × 3 filters with a stride of 1 and zero-padding (to keep the height and width unchanged), and the max pooling layers use a 2 × 2 filter to downsample and to introduce translational invariance.

### Dual input CNN architecture

For comparison, we also created a second CNN, based on (20), that represents the prevailing approach of providing the two interacting sequences as separate inputs to the network, hereafter referred to as the *dual input* model (Fig. 4). Briefly, features are extracted independently from both inputs, before being concatenated and processed by deeper layers. Both inputs have only two dimensions, the amino acid sequence as the length and a BLO-SUM50 encoding as the depth (30). The inputs are first both passed through a layer of parallel 1D convolutions (consisting of various filter sizes ranging from 1 to 9). Each differently sized convolution for the two inputs is then concatenated alongside the length dimension, before being passed to a second layer of parallel convolutions with filter size 1. This is followed by a global 1D max pooling layer, a final concatenation, a fully connected layer and a single neuron output layer. No batch normalization is used and there is only a single dropout layer, located behind the fully connected layer. Sigmoid activation functions are used throughout the network.

**Fig. 4.**
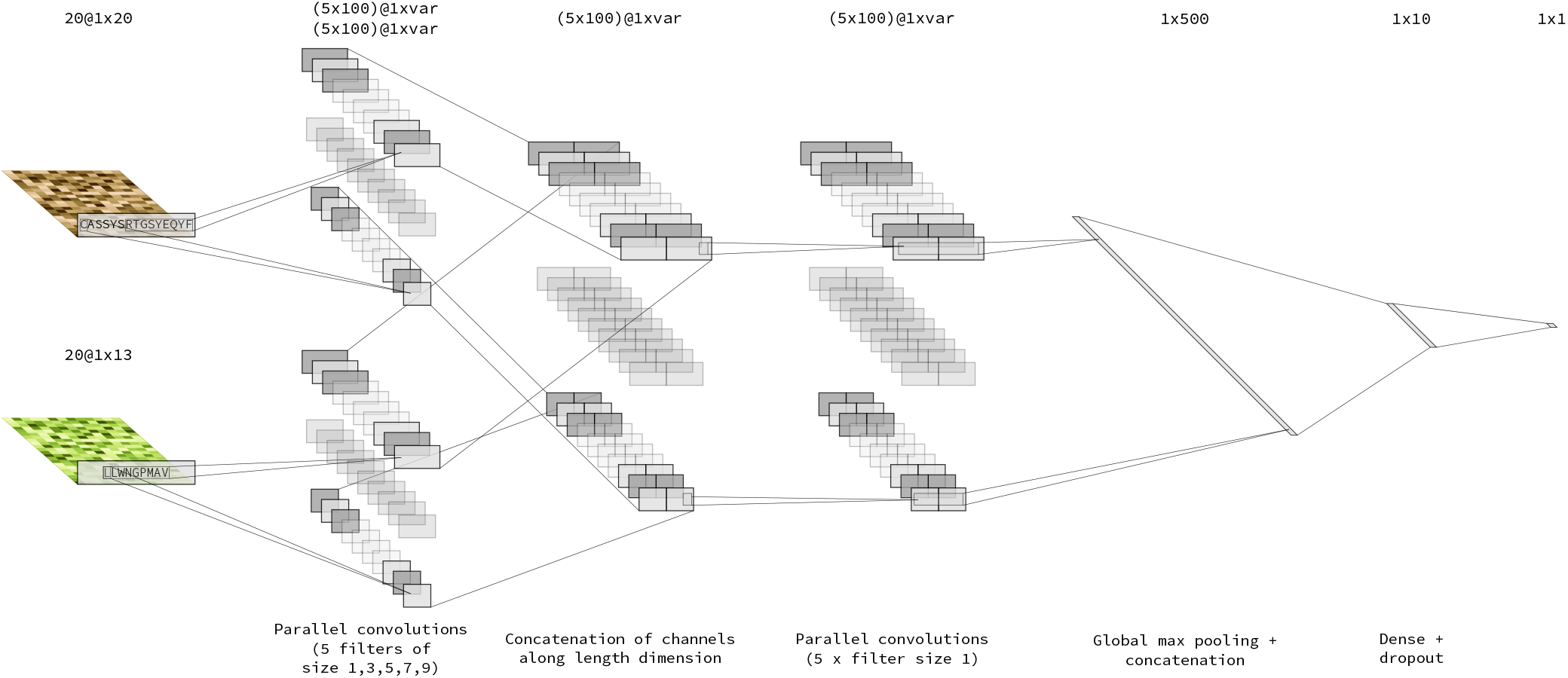
Architecture of the dual input CNN. The two input sequences are encoded using the BLOSUM50 matrix and supplied to the model in separate layers before being combined. Note that all layers contain sequences, not matrices as in the ImRex model Fig. 3, and thin black lines originating from a smaller rectangle represent 1D convolutions, whereas other lines represent concatenation steps. The length dimension of the sequences is omitted, since the differently sized kernels all result in different output lengths.

### Model training

The RMSProp optimizer (31) was used to update model weights with a learning rate of 0.0001 for the ImRex CNN and 0.001 for the separated input CNN (as in the original publication (20)). Binary cross-entropy was chosen as the objective function. The batch size was set to 32, as recommended by (32) for improved generalization performance. Models were trained for 20 epochs, which proved sufficient by inspecting loss curves.

We implemented and trained our models using TensorFlow 2.1.0 (33) and the Python (3.7.6) packages Biopython 1.76 (29), NumPy 1.18.1 (34), pandas 1.0.1 (35), pyteomics 4.2 (36, 37), scikit-learn 0.22.1 (38) and SciPy 1.5.0 (39).

### Performance evaluation for known and unknown epitopes

The CNNs were trained under two different seeded cross-validation (CV) schemes (except when explicitly mentioned otherwise): 1) a regular 5 times repeated 5-fold CV and 2) a 5-fold epitope-grouped CV. The former can be interpreted as “*predicting the recognition of known epitopes by TCRs*”, since the epitopes that are encountered during testing have already been observed during training (albeit paired with different TCR sequences). The epitope-grouped CV strategy on the other hand ensures that the TCR-epitope pairs in different folds never share their epitope (i.e. every fold contains unique epitopes). Thus, it provides a representation of the performance of the model on novel, unseen epitopes: (*epitope-agnostic* generalization performance). In order for the comparisons between different model architectures trained in this manner to be valid, the folds (and negative data generation) for experiments on the same datasets used the same seed. Generalization performance was measured using the sample mean and sample standard deviation (*s*) of the area under the receiver operating characteristic (AUROC) and average precision (AP) on held-out CV folds. Significant differences in performance were established using non-parametric testing (paired or unpaired as appropriate).

In addition to CV validation performance, the McPAS-TCR dataset (10) was used as an external test by splitting it up into two subsets: the shared epitope sub-set containing 4101 unique pairs covering 46 epitopes, which were present in the training data, and the unique epitope sub-set containing 736 pairs covering 10 unique epitopes, which were not present in the training data.

### Decoy epitope models

A challenging aspect of generic TCR-epitope interaction models is that they could potentially memorize CDR3 motifs regardless of the epitope partner. For these decoy datasets, every unique epitope was replaced by a uniform random permutation of the 20 amino acids of the same length as the original, in order to remove any chance of true molecular binding. If the performance on a decoy dataset is comparable to the real data, it seems likely that even though the model can deliver accurate predictions, it has not been able to capture the true underlying molecular forces that govern the binding process and has merely managed to memorize spurious motifs that are present within the TCR training data.

## Results

### Impact of feature encoding on known-epitope performance

To assess the known-epitope performance of the two different CNN architectures on TCR-epitope data, namely the ImRex and dual input approaches, we trained the beta chain dataset using 5 times repeated 5-fold CV with negative data generated through shuffling. The validation receiver operating characteristic (ROC) and precision-recall (PR) curves for both models are provided in Fig. 5.

**Fig. 5.**
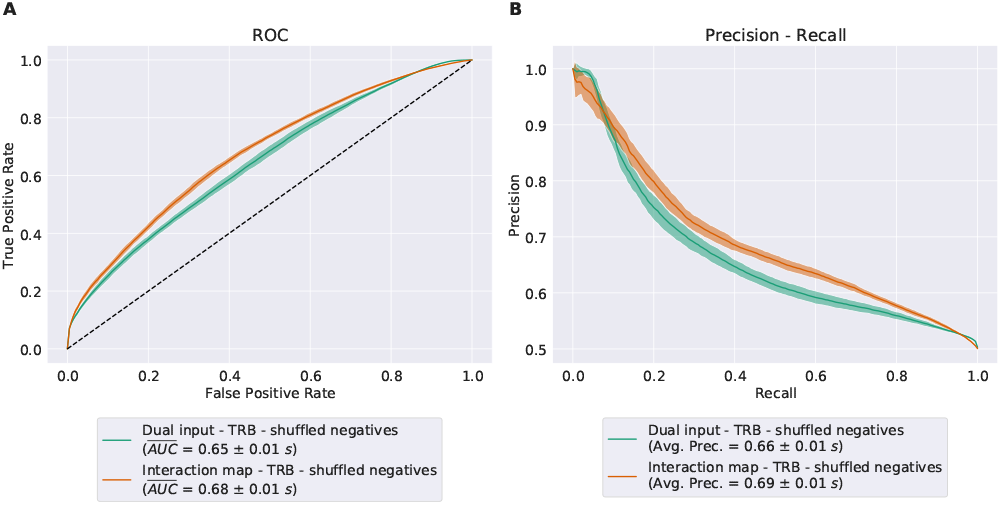
Validation ROC and PR curve for the ImRex and dual input CNNs when trained on the beta chain dataset with shuffled negatives using 5 times repeated 5-fold CV.

The ImRex model outperforms the dual input one slightly according to both AUROC (*AUROC_ImRex_* = 0.68 ±0.01*s*; *AUROC_dual_* = 0.65±0.01*s*; Mann–Whitney U-test (MWU) *P*-value =8 × 10^-10^) and AP (*AP_ImRex_* = 0.69 ± 0.01*s*; *AP_dual_* = 0.66 ± 0.01*s*; MWU *P*-value = 3.3 × 10^-9^). It is clear that for both models recall (or sensitivity) is the biggest problem; mistakes where the models fail to detect a true interaction are the most common.

The per-epitope AUROC ranking, derived by evaluating on the positive and negative TCRs for one specific epitope in each CV fold, is shown in Fig. 6.

**Fig. 6.**
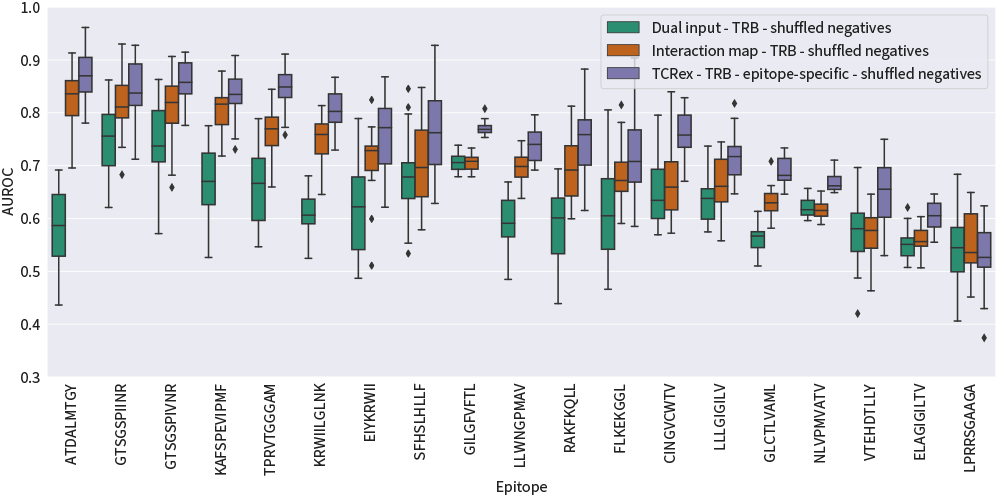
Ranking of the validation AUROC per epitope (≥ 30 examples, present in all 25 CV folds and at least one model *AUROC* > 0.50) for the dual input and ImRex CNN trained on the beta chain dataset with shuffled negatives using 5 times repeated 5-fold CV. The performance of equivalent epitope-specific RF models (TCRex) are also shown.

The differences between the two models are consistent with the overall performance, with the dual input model lagging behind the ImRex variant for most epitopes (Wilcoxon signed-rank test on per-epitope mean AUROC **P**-value = 3.8 × 10^-5^). The top epitopes perform substantially better than the overall AUROC, but it is also apparent that other epitopes do much worse. Importantly, we found no correlation between the AUROC of an epitope and the number of training examples that were available in the training set (Fig. S9). This per-epitope view also highlights that the overall performance metrics can be misleading in certain situations. Indeed, if a model works exceptionally well or poorly on a given abundant epitope, this could skew the overall performance metric in an unintended manner.

In order to compare these results to the current state-of-the-art epitope-specific classifiers, we trained equivalent random forest (RF) models for each epitope that met the above criteria based on the methodology of TCRex (19). We re-used the CV folds of the CNN models, each time extracting only those CDR3 sequences that were paired with a given epitope (both positives and negatives). Thus the TCRex performances could be overly conservative as the negative data is far more limited than that used for typical training.

As expected however, the epitope-specific models do still outperform the generic ones (Wilcoxon signed-rank test on per-epitope mean AUROC of the ImRex and TCRex models *P*-value = 1.1 × 10^-5^). Interestingly, there is a clear trend across the different models: in general, epitopes which do well, do well across all models, and vice versa, and this seems to be independent of the number of training examples.

### Sources of biases in performance

Prior to exploring the performance of various methods in the epitope-agnostic, it is worthwhile to explore the inherent challenges that the TCR-epitope recognition problem brings. In this setting, training and validation data will be used in an epitope-grouped format, which ensures that epitopes used as validation data do not occur in training data. This is a far more complex set-up than the epitope-specific case.

Various aspects of the TCR-epitope interaction problem could hinder model performance or lead to erroneous conclusions and overly optimistic (or pessimistic) estimates of generalization performance. We identified the following potential obstacles and sources of bias, and designed strategies to investigate their influence or reduce their impact during both model training and evaluation.

i. Absence of true negative data
ii. Imbalanced epitope distribution
iii. Different TCR chain inclusion
iv. Known versus unknown epitope prediction

#### Reference negatives result in over-optimistic performance compared to negatives generated via shuffling

The TCR-epitope interaction datasets only contain positive examples: interacting pairs of TCRs and epitopes. However, in order to train supervised models both positive and negative examples (pairs of TCRs and epitopes that do not recognise each other) are required. The latter need to be generated in a biologically plausible manner, as they need to represent an unbiased estimate of the true distribution of non-interacting sequences. We compared two different methods for generating negative observations: 1) shuffling the known positive pairs and 2) sampling uniformly from a negative TRB CDR3 reference repertoire set. The frequency of positive and negative examples was kept equal in both scenarios (50:50 ratio), and downsampling was used to address epitope-imbalance (see next section).

For the shuffle method, each CDR3 sequence in the positive dataset was combined with an epitope that was sampled uniformly at random from the epitopes in the positive sequence pairs, while excluding its known true peptide partner(s). This approach ensures that every CDR3 sequence occurs both in a positive and in a negative pair, only combined with a different epitope.

For the negative reference approach, CDR3 sequences sampled from the negative TRB CDR3 reference dataset were matched with epitopes sampled uniformly at random from the positive pairs. This builds on the assumption that TCRs from this reference set are unlikely to bind a randomly assigned epitope. In this case, there is no overlap between the CDR3 sequences in the positive and negative pairs, unlike in the shuffled dataset.

In isolation, it seems that the model that was trained on reference negatives does slightly better than the one trained on shuffled negatives, as can be seen in Fig. 7 (*AUROC_shuffled_* = 0.55 ± 0.02*s*; *AUROCreference* = 0.57 ± 0.03*s*). However, when they are contrasted to models trained on decoy datasets, it becomes clear that the baseline performance of reference negatives is slightly better than random, whereas we would expect the performance on the decoy dataset to be close to random (*AUROC_shuffled-decoy_* = 0.50 ± 0.01*s*; *AUROC_reference-decoy_* = 0.55 ± 0.02*s*). This indicates that the performance of the model is for the most part due to its ability to separate positive TCRs from negative TCRs regardless of the matched epitope. This may be due to an unknown bias either within the TCR-epitope data or within the reference repertoire data used for negative interactions. In contrast, when shuffled negatives are used, the decoy model has an average AUROC just below that of a random classifier, while the model trained on the true data was able to achieve a somewhat higher performance (albeit lower than the reference negative setting). This finding was also supported by a MWU, which found a significant difference between the AUROC of the true and the decoy dataset in the case of shuffled negatives (*P*-value = 0.0061), but not for the reference negatives (*P*-value = 0.11).

**Fig. 7.**
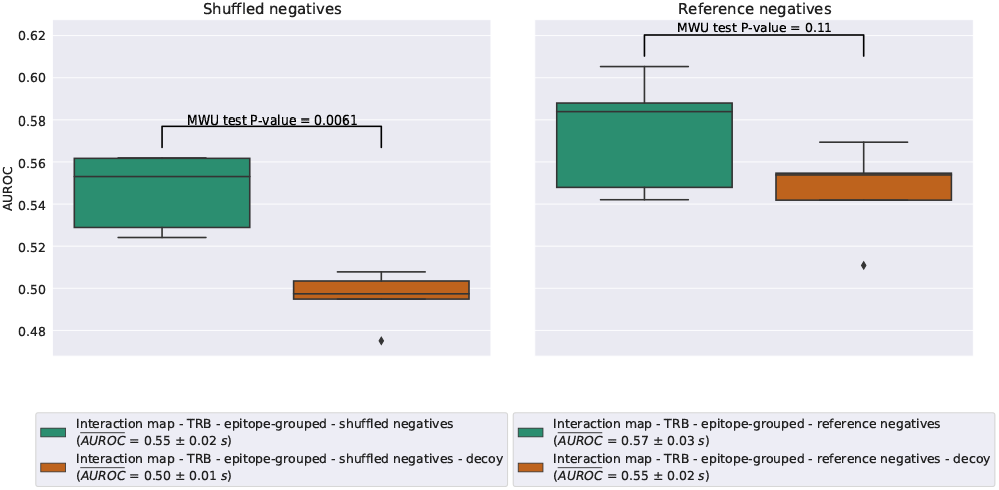
Box plots of the validation AUROC for the ImRex CNN trained on the downsampled beta chain dataset and its decoy, with either shuffled or reference negatives and using 5-fold epitope-grouped CV.

An inherent bias that comes with using an external negative reference dataset is not unexpected, and this is why prior evaluations have always used shuffled datasets. Thus for the remainder of this study, we will continue with only the shuffled negative dataset.

For the known-epitope models the differences between true and decoy data very small, but significant for both types of negatives, see Fig. S1.

#### Downsampling to address epitope imbalance has a small negative effect on model performance, but allows for more robust CV strategies

As can be seen from Fig. 2, the number of interaction pairs differs substantially between different epitopes. This has two important implications for evaluating the performance of our predictive models.

First, models could develop a bias towards (i.e. overfit on) those epitopes which deliver the majority of the training data. To combat this, model performance was not only evaluated on the entire test dataset (which results in an overall metric), but also on an individual per-epitope basis. A per-epitope metric can immediately reveal whether a model has focused exclusively on a small number of abundant epitopes, and at the same time ensures that the performance evaluation is not steered entirely by the most abundant epitopes, unlike the overall performance which is strongly influenced by those. This entails evaluating the trained model on a test set (or its held-out CV folds) that is restricted to only those sequence pairs that contain a specific epitope (both positives and negatives).

Second, when an epitope-grouped CV strategy is used, it might not be possible to create folds that are balanced in terms of both the total number of interactions and the number of unique epitopes. Folds might even be created that only contain a single abundant epitope, making it impossible to generate negative pairs via shuffling. To avoid this issue, a downsampling strategy was employed, where the number of interactions for the top outliers was reduced through uniform sampling. While this does not entirely fix the imbalance, it does lower the highest counts to less than the CV fold size, ensuring that a mix of epitopes is present in each fold. In short, downsampling allowed us to create epitope-grouped CV folds with a more comparable epitope distribution, at the cost of having substantial fewer observations for training and testing.

To gauge the effects of this downsampling strategy, models were trained using either moderate or strong downsampling, by setting a threshold of 1,000 and 400 examples per epitope respectively (reducing the beta chai dataset from 14,188 to 8,945 and 6,702 sequence pairs respectively), or no downsampling at all (see Table S1 for which epitopes were impacted). No downsampling restricted the number of possible epitope-grouped CV folds to three instead of five when using shuffled negatives. These scenarios examine the trade-off between sample size and epitope-skew, and at the most extreme end (i.e. no downsampling) combines this with a less robust estimate of model stability due to the lower number of CV folds. As shown in Fig. 8, the differences in AUROC between the different degrees of downsampling (*AUROC_no_* = 0.58 ± 0.04*s*; *AUROC_moderate_* = 0.54 ± 0.04*s*; *AUROC_strong_* = 0.55 ± 0.02*s*) are not significantly different according to the Kruskal-Wallis test (*P*-value = 0.27) for the ImRex CNN model trained on the beta chain dataset with shuffled negatives.

**Fig. 8.**
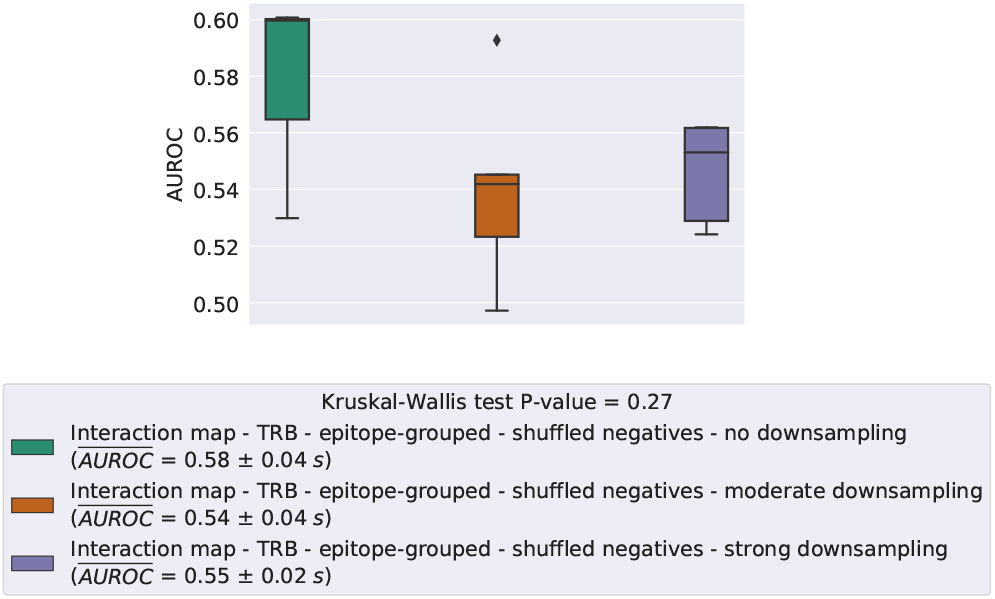
Box plots of the validation AUROC for the ImRex CNN trained on the beta chain datasets with shuffled negatives and epitope-grouped CV using different degrees of downsampling: none (3-fold CV), moderate downsampling with a threshold of 1000 examples per epitope (5-fold CV) and strong downsampling with athreshold of 400 examples per epitope (5-fold CV).

Importantly, the per-epitope performance of the most abundant epitopes (e.g. NLVPMVATV and GILGFVFTL) are similar among all three datasets (Fig. S2). Consequently, the observed differences in performances can likely be attributed mostly to differences in CV splits and negative data generation, and the larger training size when not downsampling and using fewer folds.

These results suggest that the effect of downsampling, which is utilised throughout for the epitope-agnostic setting, will not dominate the effects on performance. Because we focus on per-epitope comparisons in this study, we opted to use the strong downsampling variant since it allows for a more thorough CV strategy and because it is least problematic in terms of maintaining a balanced ratio of positives and negatives on a per-epitope basis.

The effects of downsampling appear to be negligible for the known-epitope models, as shown in Fig. S3.

#### TCR chain information is not exchangeable

While TCRs are made up of two chains, the TRB chain is far more ubiquitous in TCR-epitope databases. A naive approach for modeling recognition may be to utilize both chains as individual examples, since similar molecular interaction patterns should underlie both chains due to the fact that they are similar in composition and that they both contact the same molecule, i.e. the epitope.

The models that were trained on the beta chain dataset outperform those that were trained on the mixed chain dataset (*AUROC_mixed_* = 0.52 ± 0.01*s*; *AUROCb_eta_* = 0.55±0.02*s*, MWU test *P*-value = 0.030, Figures S4 and S5). In contrast, models that were trained purely on TRB data do not generalize to the alpha chain dataset, and the performance is no better than random (*AUROC_alpha_* = 0.47).

Similar observations hold for the known-epitope setting (see Figures S6 and S7), models trained on exclusively TRB sequences do not carry over to TRA sequences. However, here the mixed chain dataset does lead to better results than the beta chain dataset.

### Impact of feature encoding on epitope-agnostic performance

Bearing in mind the issues that were discussed in the previous section, we compared the generalization capabilities of the two CNN architectures to unseen epitopes via an epitope-grouped CV on the beta chain dataset while using strong downsampling and negatives generated by shuffling the sequence pairs.

Both the ImRex and dual input CNN show a higher than random, but generally poor epitope-agnostic performance (Fig. S8): *AUROC_ImRex_* = 0.55±0.02*s*, *AUROC_dual_* = 0.52 ± 0.01*s* (MWU *P*-value = 0.018) and *AP_ImRex_* = 0.55 ± 0.02*s*, *AP_dual_* = 0.52 ± 0.01*s* (MWU *P*-value = 0.018). Thus as expected the performance is far worse for the epitope-agnostic problem than for the known-epitope problem that was shown before. This signifies that the performance on known epitopes cannot simply be transfered to this more challenging problem.

Importantly, however, a ranking of the per-epitope AUROC shows that despite the lacklustre overall performance, the models do manage to perform reasonably well for a small subset of epitopes (Fig. 9). The top ranking individual epitope performances are achieved by the ImRex model (eight out of the ten highest AUROC values). This is confirmed by a Wilcoxon signed-rank test, which finds a significant difference (*α* = 0.05) in the per-epitope AUROC medians (*P*-value = 0.016), although naturally this test depends strongly on the selected thresholds for the AUROC and number of examples. These results once again highlight the fact that overall performance metrics cannot provide a complete picture of the behaviour of TCR-epitope prediction models.

**Fig. 9.**
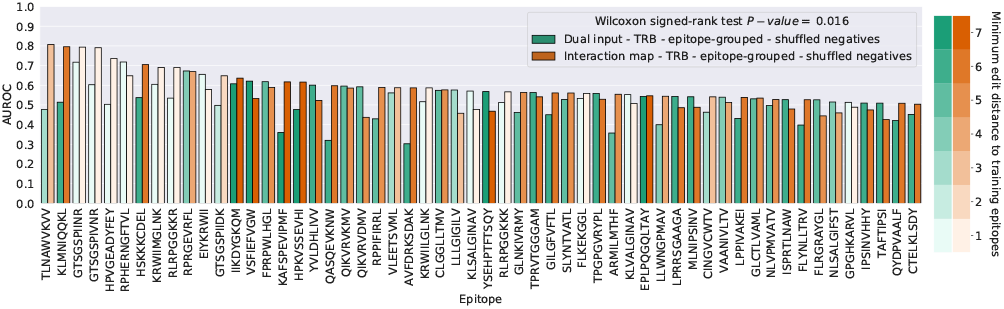
Ranking of the validation AUROC per epitope (≥ 30 examples and at least one model *AUROC* > 0.50) for the ImRex and dual input CNNs after 5-fold epitope-grouped CV when trained on the downsampled beta chain dataset with shuffled negatives. The color gradient indicates the minimum edit distance of each epitope to the epitopes in the training dataset.

#### Strong performance for closely related epitopes

When the epitopes that show a strong performance for the ImRex approach are compared with the epitopes that were present in the training dataset, a striking pattern emerges. As can be seen in Fig. 10, the highest performance is achieved for those epitopes that shared some similarity with epitopes present in the training data. For epitopes with at most a single amino acid difference, the average AUROC over the epitopes is 0.61, which is significantly higher than the average decoy AUROC of 0.49 (MWU test *P*-value = 0.0004). Thus the TCR-epitope patterns learned by the molecular ImRex approach, work best for related epitopes. This is not unexpected as these epitopes share similar physicochemical properties and thus some underlying interaction patterns may be shared. It is then the task of the CNN to learn which patterns are relevant in which case. In contrast, the dual input approach was unable to achieve this relationship (mean AUROC for epitopes with minimum edit distance 1 = 0.51; mean AUROC for decoy epitopes = 0.50; MWU test *P*-value = 0.28), as can be seen in Fig. S10, demonstrating that this is a non-trivial problem.

**Fig. 10.**
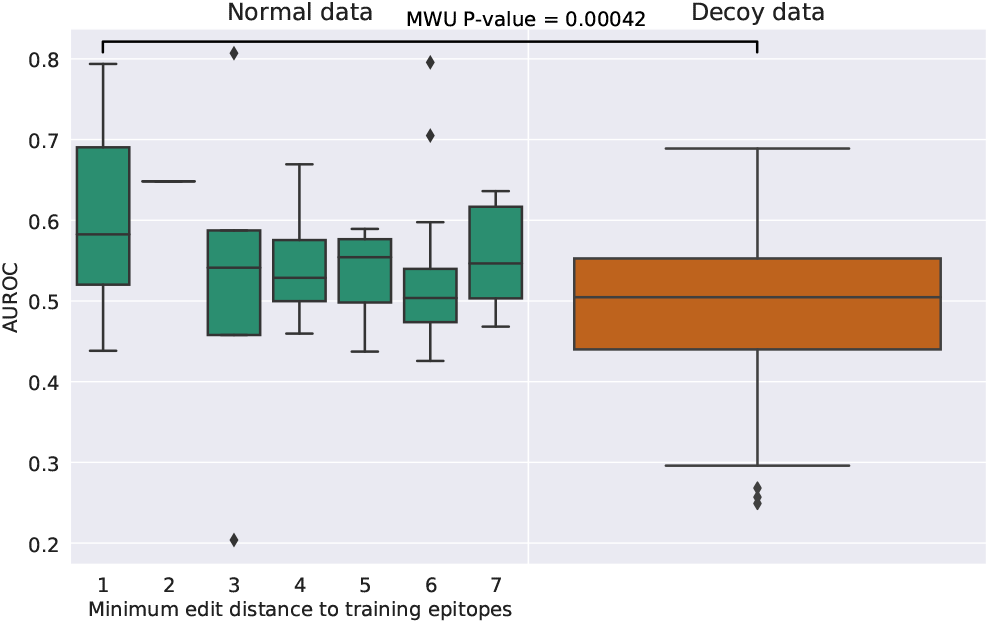
Box plots of per-epitope AUROC and minimum edit distance to epitopes in the training dataset for the ImRex CNNs trained using 5 times repeated 5-fold CV on the downsampled beta chain dataset with shuffled negatives.

These findings also highlight that the overall AUROC achieved by a model can be misleading in certain situations, as in this case, where the average AUROC over the different CV folds was influenced by a small number of outlying epitope examples (each occurring fewer than 30 times). Depending on one’s perspective, either the overall or the per-epitope performance is the more appropriate measure of performance, but a distinction must be made regardless.

### Evaluation on independent datasets

We utilised the McPAS-TCR dataset (10) as an independent test set. After filtering, it was split into two subsets: one containing epitopes already present in the VDJdb training data, and one with unique epitopes. As before, negative interaction pairs were generated using a shuffling approach for each subset. Both test datasets were used to evaluate the ImRex CNN that was trained on the entire downsampled beta chain dataset with shuffled negatives.

The ImRex model did not attain the same level of performance on the shared-epitope McPAS-TCR data as it did during known-epitope CV (*AUROC* = 0.59 and *AP* = 0.59). However, as before the overall performance can be considered a misleading metric here. The most abundant epitopes (LPRRSGAAGA) makes up over 35% of the observations, while showing an AUROC of 0.50, and this strongly influences the overall performance values. Looking at the the individual per-epitope performances, these seem in line with those seen during the 5 times repeated 5-fold CV, both in terms of absolute numbers and which specific epitopes do well (Figures 6 and 11).

**Fig. 11.**
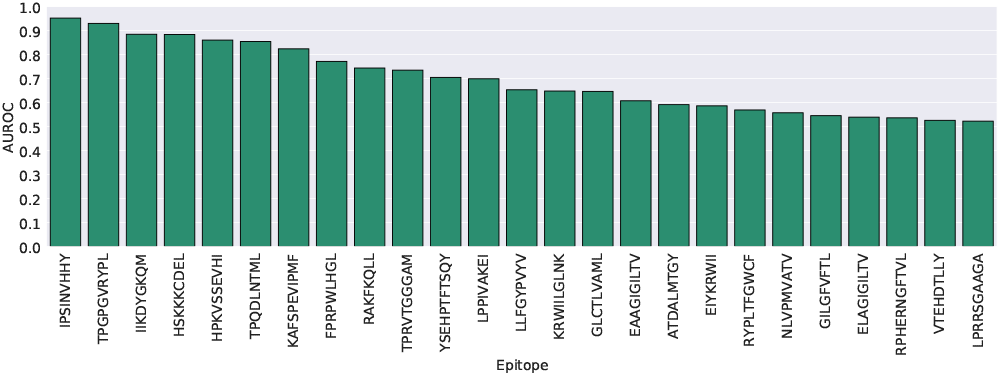
Ranking ofthe test AUROC per epitope (≥ 30 examples) on the shared epitope McPAS-TCR data for the ImRex CNN when trained on the entire downsampled beta chain dataset with shuffled negatives.

Similar to the epitope-grouped CV setting, the overall AUROC and AP were only slightly better than random performance for the unique-epitope McPAS-TCR data (*AUROC* = 0.54 and *AP* = 0.53). Bearing in mind the drawbacks of the overall performance metrics, the ranking of the individual per-epitope AUROCs are shown in Fig. 12. Like in the previous results, strong performance is found for epitopes with low edit distance.

**Fig. 12.**
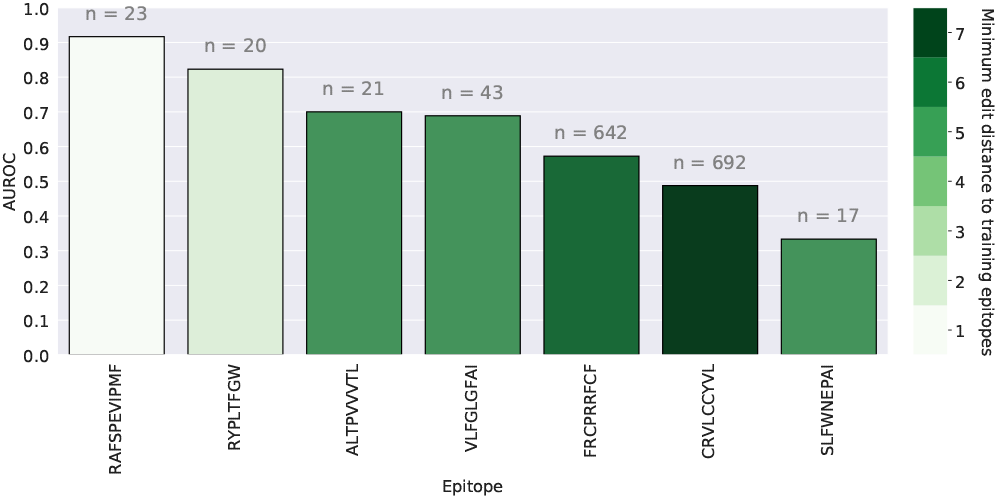
Ranking of the test AUROC per epitope (≥ 10 examples) on the unique epitope McPAS-TCR dataforthe ImRex CNN when trained on the entire downsampled beta chain dataset with shuffled negatives. The color gradient indicates the minimum edit distance of each epitope to the epitopes in the training dataset. The total number of testing examples is depicted above each bar.

## Discussion

The prediction of TCRs binding to unseen epitopes is extremely challenging. The lack of training examples, or conversely, the immense space covered by the human immune repertoire and the potentially interacting epitopes, remains an issue for the prediction of novel epitopes that were not encountered during training, and it will likely not be overcome via new modelling approaches alone. As long as the available training data only sparsely covers the epitope sequence space, it will likely limit the predictive power of any epitope-agnostic approach.

In this work, we have presented a new feature representation technique for generic sequenced-based TCR-epitope interaction prediction models, based on the pairwise combination of amino acids, called ImRex. While the overall performance remains poor, a comparison with a standard dual input embedding approach showed an improved performance of the ImRex model for the epitope-agnostic setting. This is due in part to a high performance unique to ImRex for epitopes which have similar amino acid patterns to the training data. This supports our hypothesis that the use of an interaction map, instead of separate embedding steps for the two molecular interactors, allows the model to identify molecular TCR-epitope interactions which may have helped to avoid overfitting and improve the generalization capabilities of the model.

A similar increase in performance of the ImRex model compared to the dual input approach could be found for the known-epitope setting. However, here our results show that these models also display high performance trained on decoy data with nonsense epitopes. We hypothesize that the current generation of epitope-agnostic still mainly rely more on CDR3 patterns for the prediction of TCR interactions with known epitopes. Hence, they work similar to epitope-specific models, such as TCRex, for known epitopes. Thus the behavior for the known epitope setting cannot be readily extrapolated to that of the epitope-agnostic setting. In the case of ImRex, we do find a minor increase above the decoy performance, indicating that it may still be using the epitope information itself in a small part for known epitopes.

We have also shown that overall performance metrics might not be the most appropriate measure to assess generic TCR-epitope interaction prediction models, as this metric could be strongly influenced by the abundance of specific epitopes with better (or worse) than average performance.

Furthermore, we listed a number of other challenges that are unique to the TCR-epitope prediction problem, and examined different approaches to deal with them. These were related to the type of negative data generation, the epitope imbalance and the exchangeability of TCR alpha and beta chains. We also demonstrate the use of decoy epitope data as a method of establishing a baseline performance.

First, we recommend the use of shuffling over a reference repertoire as the de-facto method of generating negatives in most situations. In the epitope-agnostic case only shuffled negatives outperform their decoy counterpart (albeit at a lower overall performance than reference negatives). For the known-epitope setting the impact on model performance seems to be fairly limited however.

Next, no clear trends were found between the number of training examples of a given epitope and its final individual performance in the known-epitope setting. This might be explained by the underlying diversity of the epitope-specific TCRs. In general, we expect to overestimate the model performance when the model is trained on a low-diversity data set which does not give a good representation of the entire epitope-specific repertoire (i.e. few large TCR clusters). Ideally future evaluation metrics and training procedures should attempt to take this diversity into account. Although, the size of the training dataset did not seem to impact the model performance, downsampling will remain necessary when an epitope-grouped CV is required, depending on the desired number of folds and the extent of the epitope-skew. The fact that the amount of downsampling has a negligible effect on the epitope-grouped models supports our assumption that future improvements to TCR-epitope recognition prediction will largely depend on the collection of more diverse epitope-TCR pairs.

Finally, we found that TCR chain information is not exchangeable. In other words, adding TRA chain sequences as additional training examples does not markedly improve model performance, nor can models trained on TRB chains be transferred to datasets with only TRA chains. Note that this does not preclude the possibility that models trained on a combined input of the TRA and TRB chains could improve our current results, which has been suggested by other studies (15, 40), although this does require adequate amounts of paired data.

Given these results, we believe there is a strong need for a more consistent and stringent benchmarking framework for assessing the generalization performance of TCR-epitope recognition models. The influence of epitope bias is clearly shown by the difficulty of extrapolating trained models to unseen epitopes, and as mentioned TCR diversity could have an effect here as well. A quantification method that can measure the extent of overlap between training and test data could play an important role in adressing these issues.

Lastly, it has not escaped our notice that interaction maps could be used to model other types of molecular interactions, such as protein-protein or antibody-antigen interactions.

## Key Points

1. We present ImRex, a novel feature representation method for generic TCR-epitope interaction models which combines the physicochemical properties of both interactors on the amino-acid level.
2. We demonstrate that ImRex can enhance model performance compared to separate embedding strategies, when prediction is limited to known epitopes. However, epitope-specific models still outperform generic models in this task. Epitope-agnostic prediction (based on an epitope-grouped CV) remains difficult, but we find evidence that ImRex can more easily extrapolate to epitopes that are close to the training data than dual input approaches, and achieve performance on-par with the known-epitope setting for these epitopes.
3. Overall performance metrics for generic TCR-epitope interaction models should be interpreted with care, as the performance for the individual epitopes can deviate and have a large impact on the average performance depending on the number of examples for each epitope.
4. The generation of a negative training data set through shuffling can limit performance overestimation due to memorization of CDR3 sequences.
5. The performance of epitope-agnostic models is currently still limited by the lack of sufficient amounts of diverse TCR-epitope pairs, but appropriate modelling approaches can alleviate this to a small degree.

## DATA AVAILABILITY

The code underlying this article (including a script to perform new predictions) is available on GitHub, at https://github.com/pmoris/ImRex. The data (processed train/evaluation data, trained model files, learning curves and other metrics) are available in Zenodo, at https://doi.org/10.5281/zenodo.3973547. The underlying datasets were derived from sources in the public domain: VDJdb (8) (https://vdjdb.cdr3.net/), McPAS-TCR (10) (http://friedmanlab.weizmann.ac.il/McPAS-TCR/) and TCR repertoires collected by Dean et al. (28) (https://doi.org/10.21417/B7CC7F).

## ACKNOWLEDGEMENTS

We thank Danh Bui for all the insightful discussions. We thank all contributors to VDJdb and McPAS-TCR for making their data publicly available.

## FUNDING

This work was supported by the Research Foundation – Flanders (FWO) [1141217N to P.Mo., 1S48819N to S.G., postdoctoral grant to W.B., 1861219N to B.O.], the University of Antwerp [BOF GOA: PIMS] and the Flemish Government un-derthe “Onderzoeksprogramma Artificiële Intelligentie (AI) Vlaanderen” programme. The computational resources and services used in this work were provided by the HPC core facility CalcUA of the Universiteit Antwerpen, and VSC (Flemish Supercomputer Center), funded by the Research Foundation - Flanders (FWO) and the Flemish Government – department EWI.

## COMPETING FINANCIAL INTERESTS

The authors declare no conflict of interest.

## Supplementary information

**Table S1.** Top 10 most abundant epitopes in the mixed, beta and alpha chain datasets, before downsampling.

| Mixed chain |  | Beta chain |  | Alpha chain |  |
| --- | --- | --- | --- | --- | --- |
| Epitope | Count | Epitope | Count | Epitope | Count |
| NLVPMVATV | 6452 | NLVPMVATV | 4387 | GILGFVFTL | 2433 |
| GILGFVFTL | 5289 | GILGFVFTL | 2856 | NLVPMVATV | 2065 |
| ELAGIGILTV | 1002 | ELAGIGILTV | 960 | LLLGIGILV | 330 |
| GLCTLVAML | 991 | GLCTLVAML | 883 | LLWNGPMAV | 245 |
| LLWNGPMAV | 649 | LLWNGPMAV | 404 | GLCTLVAML | 108 |
| LLLGIGILV | 561 | KRWIILGLNK | 316 | CINGVCWTV | 69 |
| KRWIILGLNK | 322 | LLLGIGILV | 231 | NEGVKAAW | 48 |
| CINGVCWTV | 199 | VTEHDTLLY | 199 | KLVALGINAV | 43 |
| VTEHDTLLY | 199 | TPRVTGGGAM | 191 | ELAGIGILTV | 42 |
| TPRVTGGGAM | 197 | RAKFKQLL | 172 | KLSALGINAV | 25 |

**Fig. S1.**
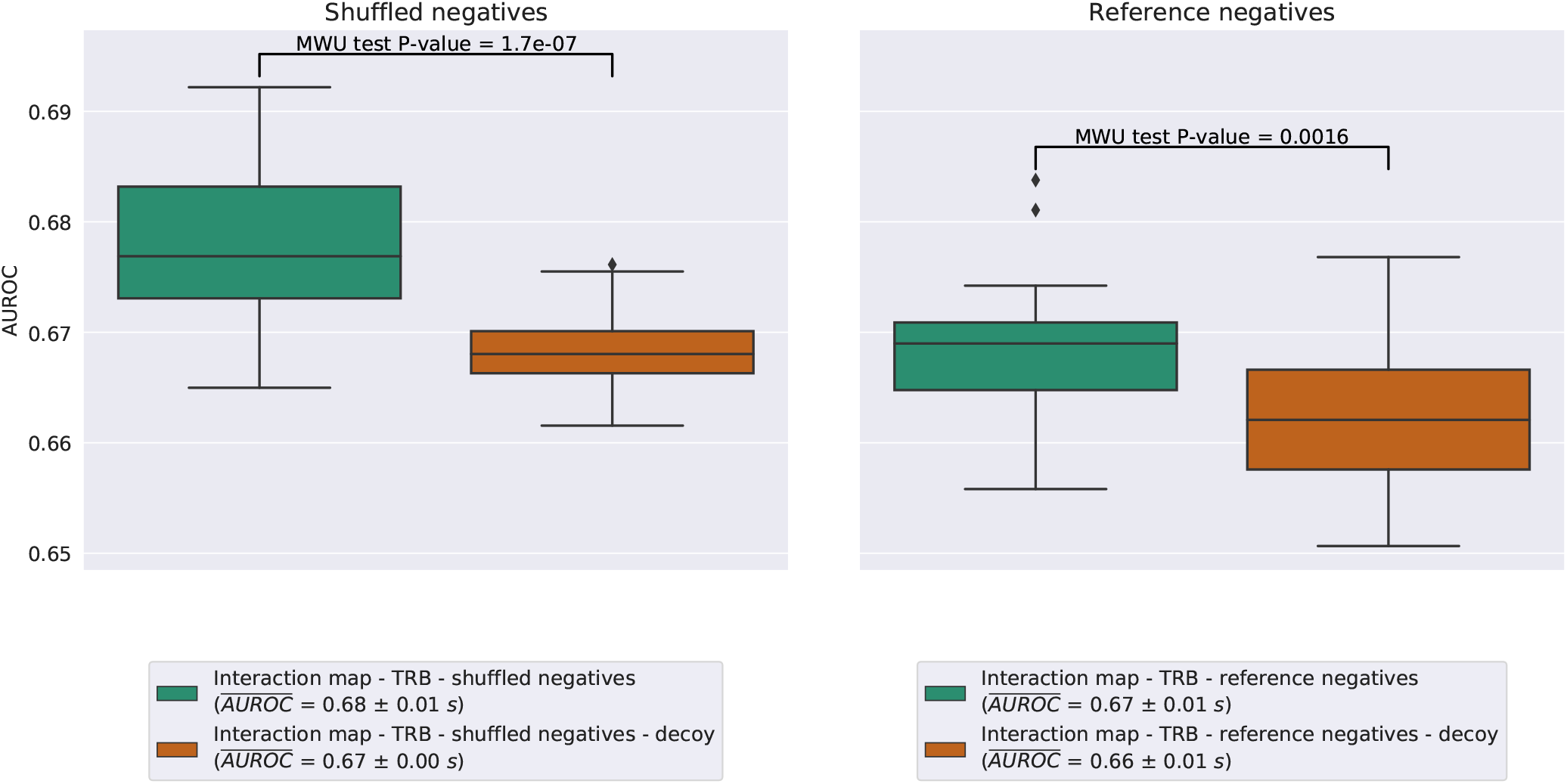
Box plots of the validation AUROC for the ImRex CNN trained on the beta chain dataset and its decoy, with either shuffled or reference negatives and using a 5 times repeated 5-fold CV.

**Fig. S2.**
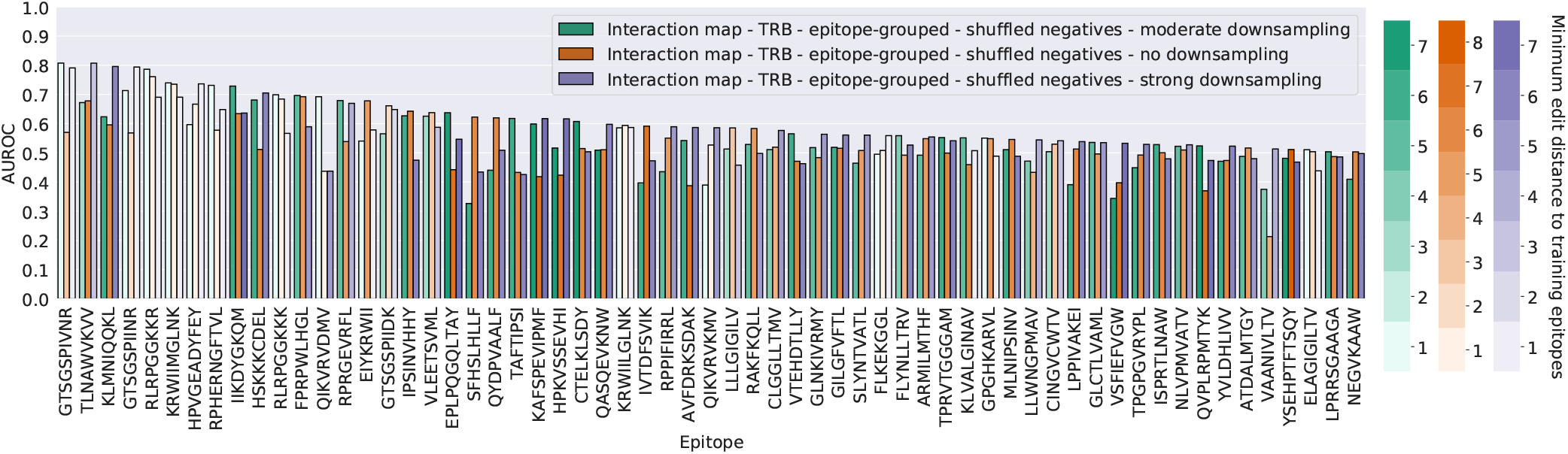
Ranking of the validation AUROC per epitope (≥ 30 examples and at least one model *AUROC* > 0.50) for the ImRex CNN trained with shuffled negatives and epitope-grouped CV using different degrees of downsampling: none (3-fold CV), moderate downsampling with a thresholdof 1000 examples per epitope (5-fold CV) and strong downsampling with a threshold of 400 examples per epitope (5-fold CV).

**Fig. S3.**
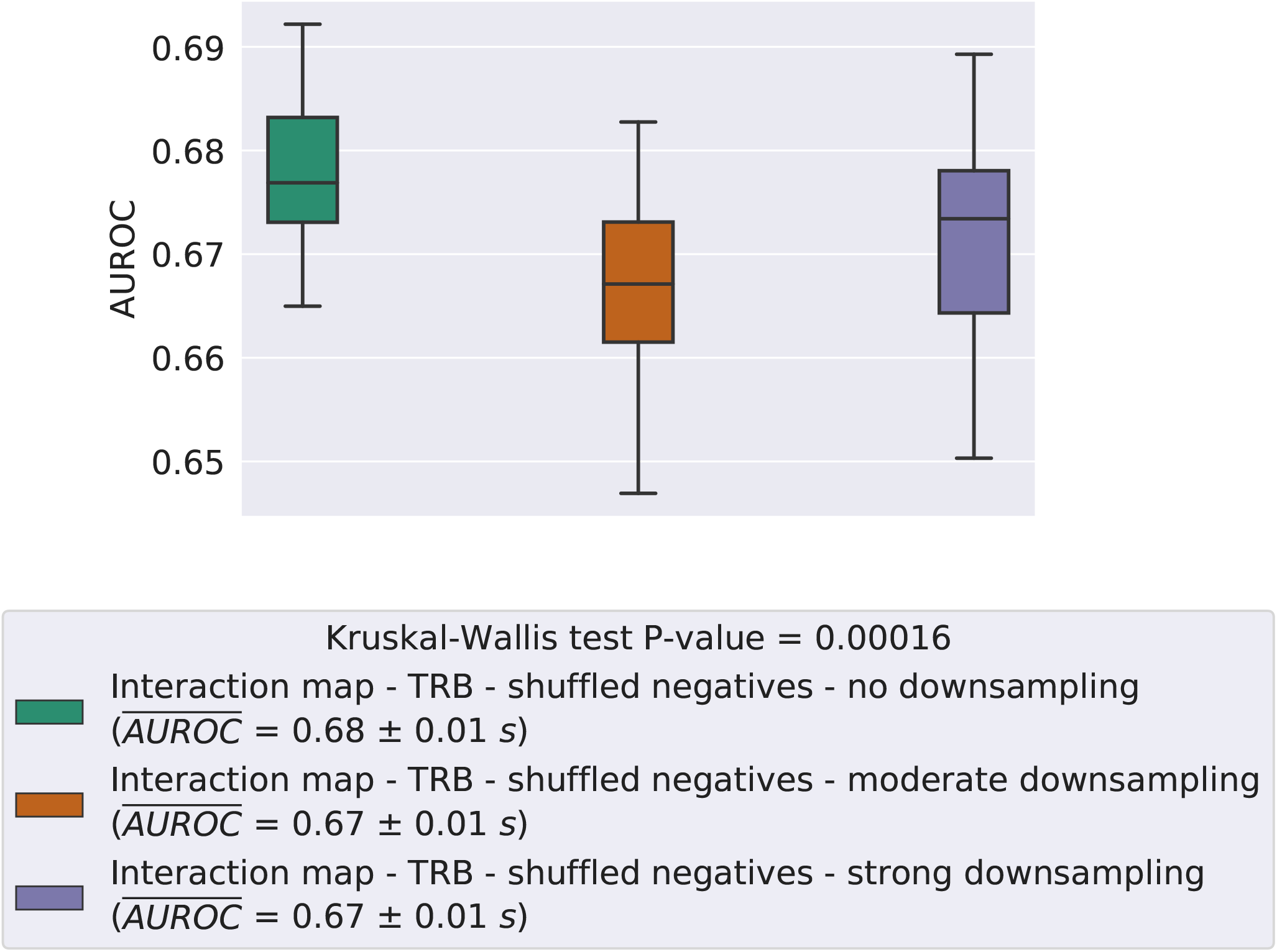
Box plots of the validation AUROC for the ImRex CNN trained on the beta chain datasets with shuffled negatives and 5 times repeated 5-fold CV using different degrees of downsampling: no downsampling, moderate downsampling with a threshold of 1000 examples per epitope and strong downsampling with a threshold of 400 examples per epitope.

**Fig. S4.**
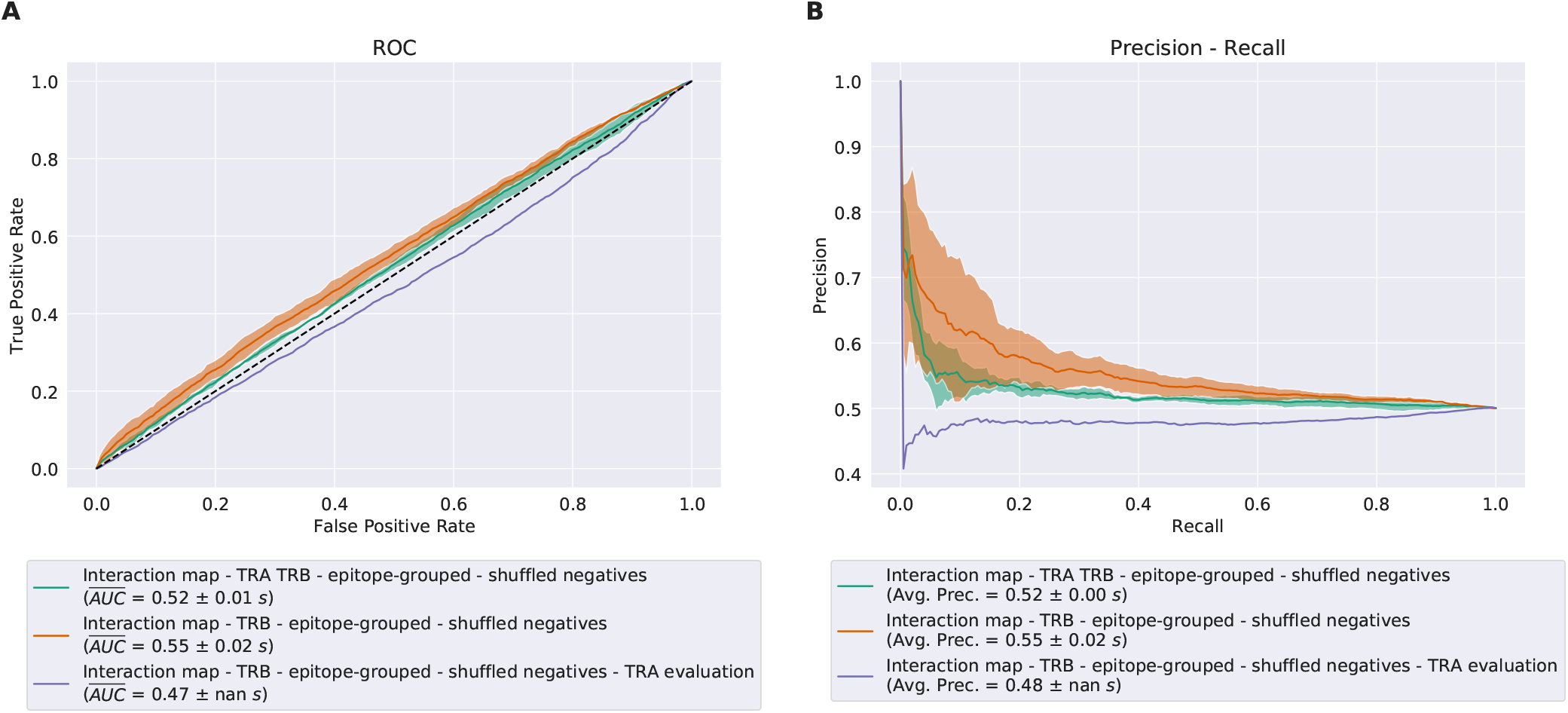
Validation ROC and PR curve for the ImRex CNN trained with shuffled negatives on the 1) downsampled mixed chain dataset and 2) downsampled beta chain dataset using 5-fold epitope-grouped CV, or on 3) the entire downsampled beta chain dataset and evaluated on the alpha chain dataset.

**Fig. S5.**
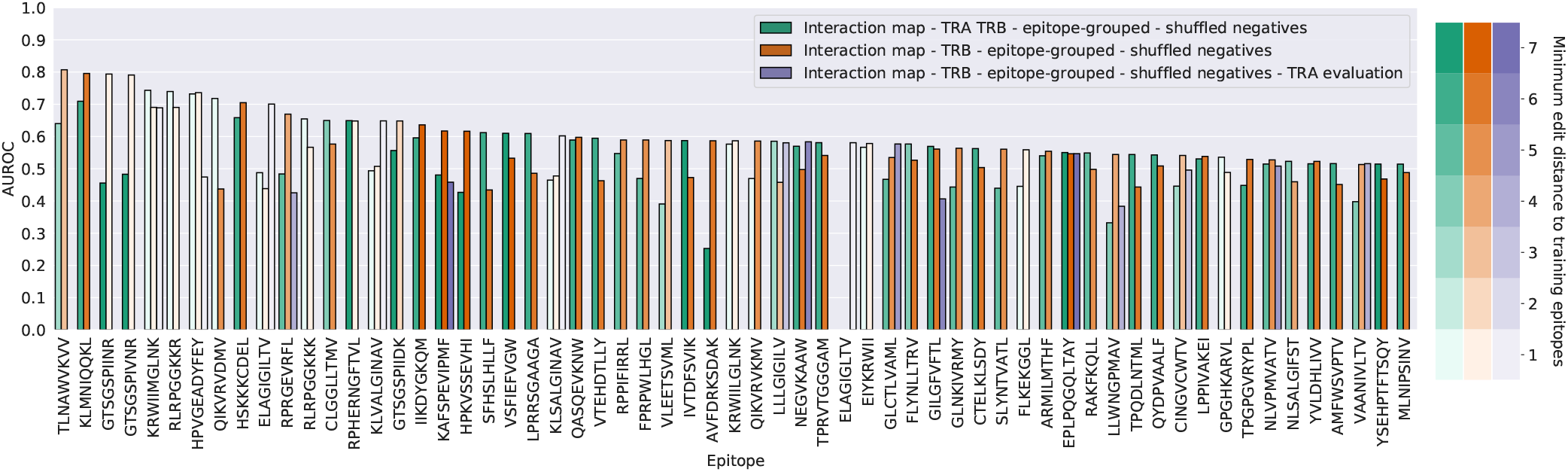
Ranking of the validation AUROC per epitope (≥ 30 examples and at least one model *AUROC* > 0.50) for the ImRex CNN trained with shuffled negatives on the 1) downsampled mixed chain dataset and 2) downsampled beta chain dataset using 5-fold epitope-grouped CV, or on 3) the entire downsampled beta chain dataset and evaluated on the alpha chain dataset.

**Fig. S6.**
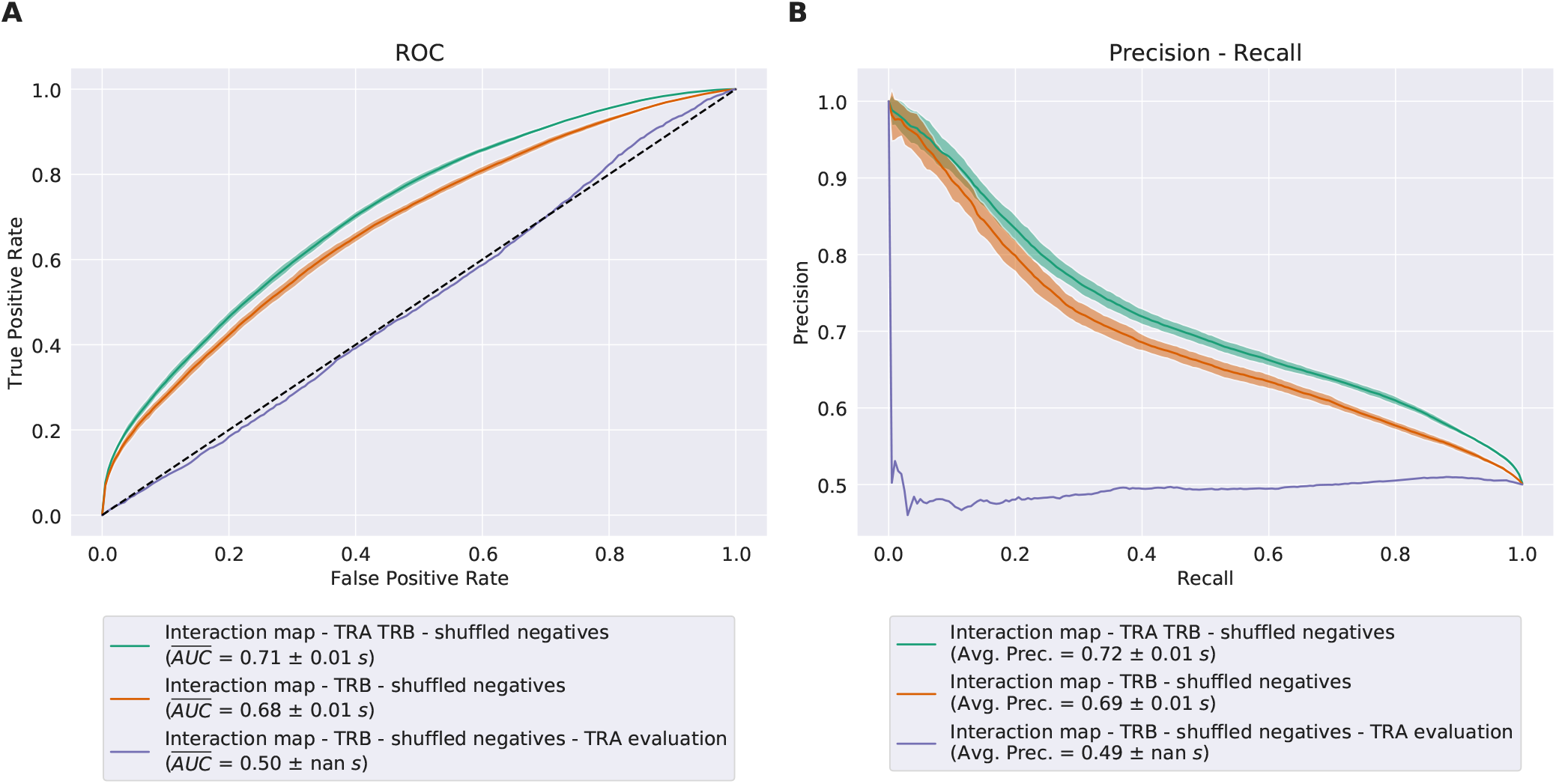
Validation ROC and PR curve for the ImRex CNN trained with shuffled negatives on the 1) downsampled mixed chain dataset and 2) downsampled beta chain dataset using 5 times repeated 5-fold CV, or on 3) the entire downsampled beta chain dataset and evaluated on the alpha chain dataset.

**Fig. S7.**
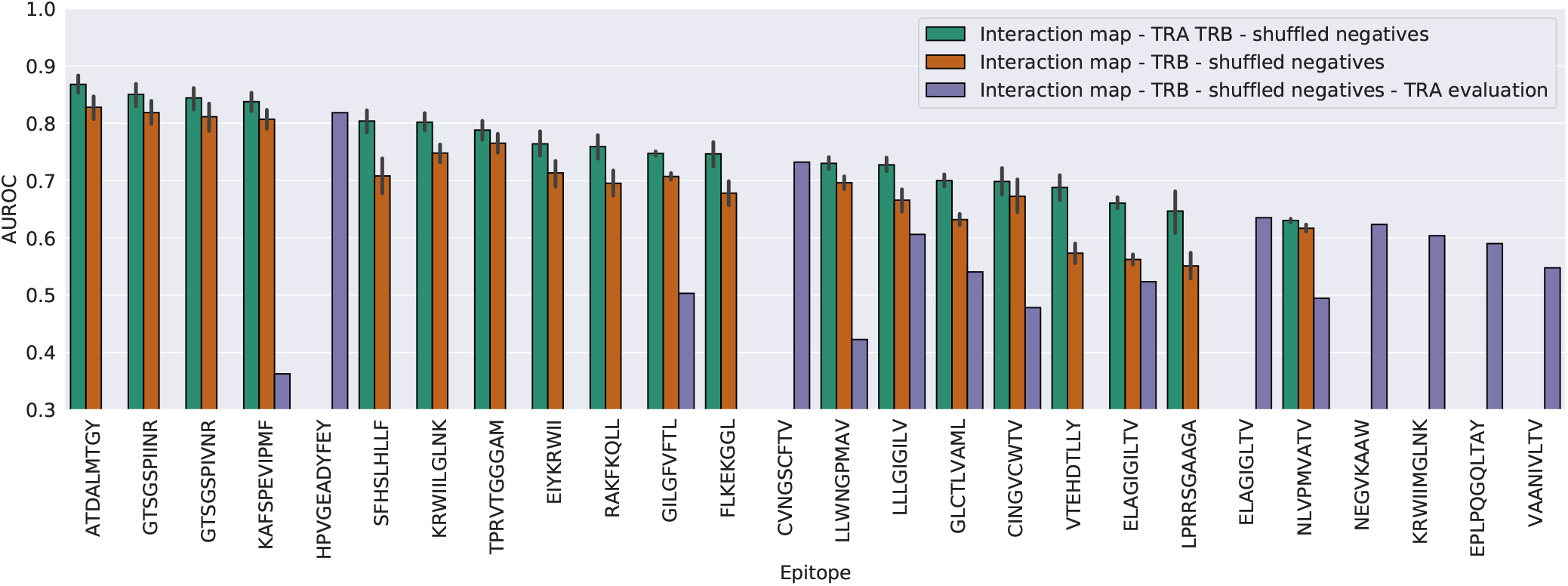
Ranking of the validation AUROC per epitope (≥ 30 examples and at least one model *AUROC* > 0.50) for the ImRex CNN trained with shuffled negatives on the 1) downsampled mixed chain dataset and 2) downsampled beta chain dataset using 5 times repeated 5-fold CV, or on 3) the entire downsampled beta chain dataset and evaluated on the alpha chain dataset.

**Fig. S8.**
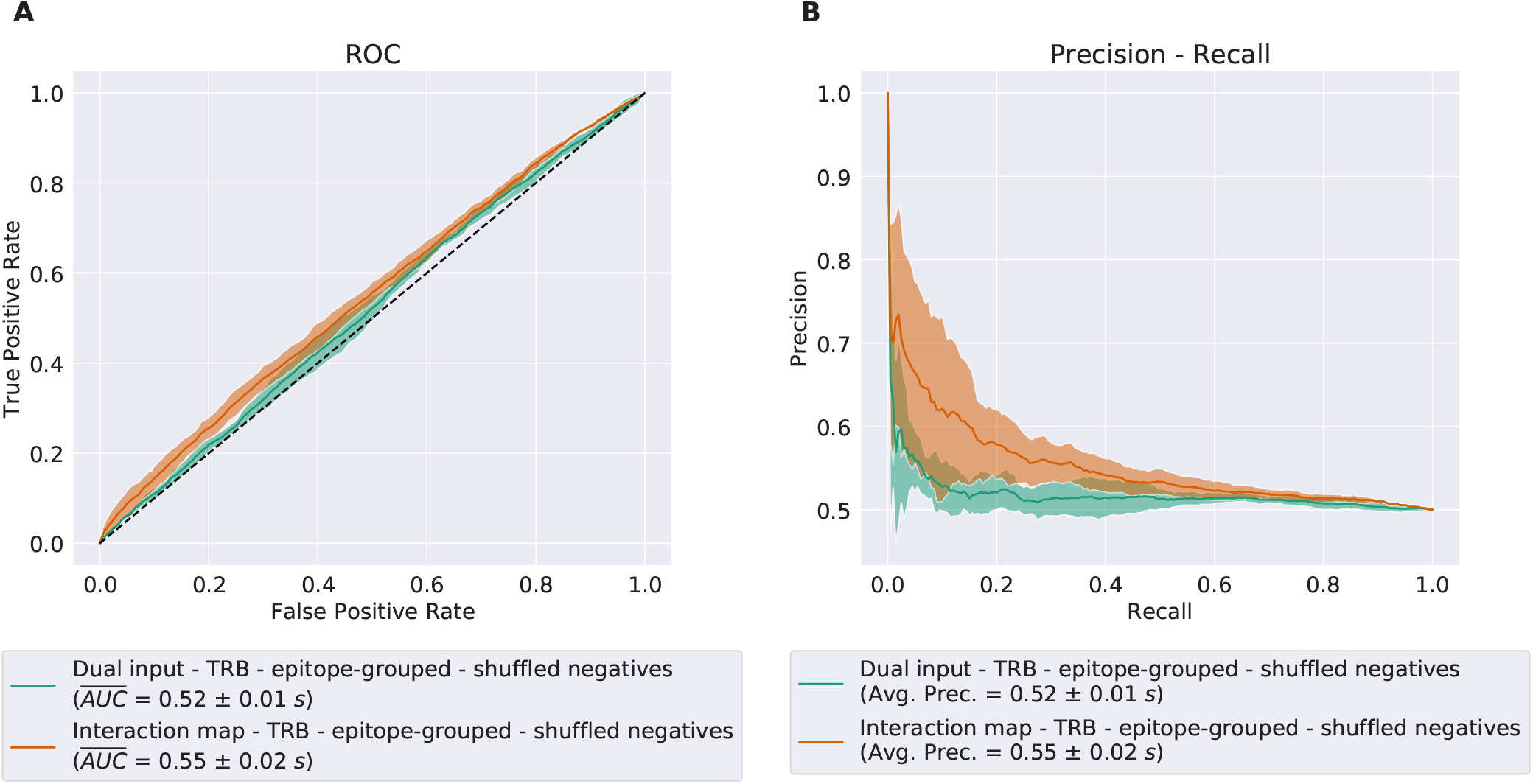
Validation ROC and PR curves for the interaction map and dual input CNN after 5-fold epitope-grouped CV, when trained on the downsampled beta chain dataset using shuffled negatives.

**Fig. S9.**
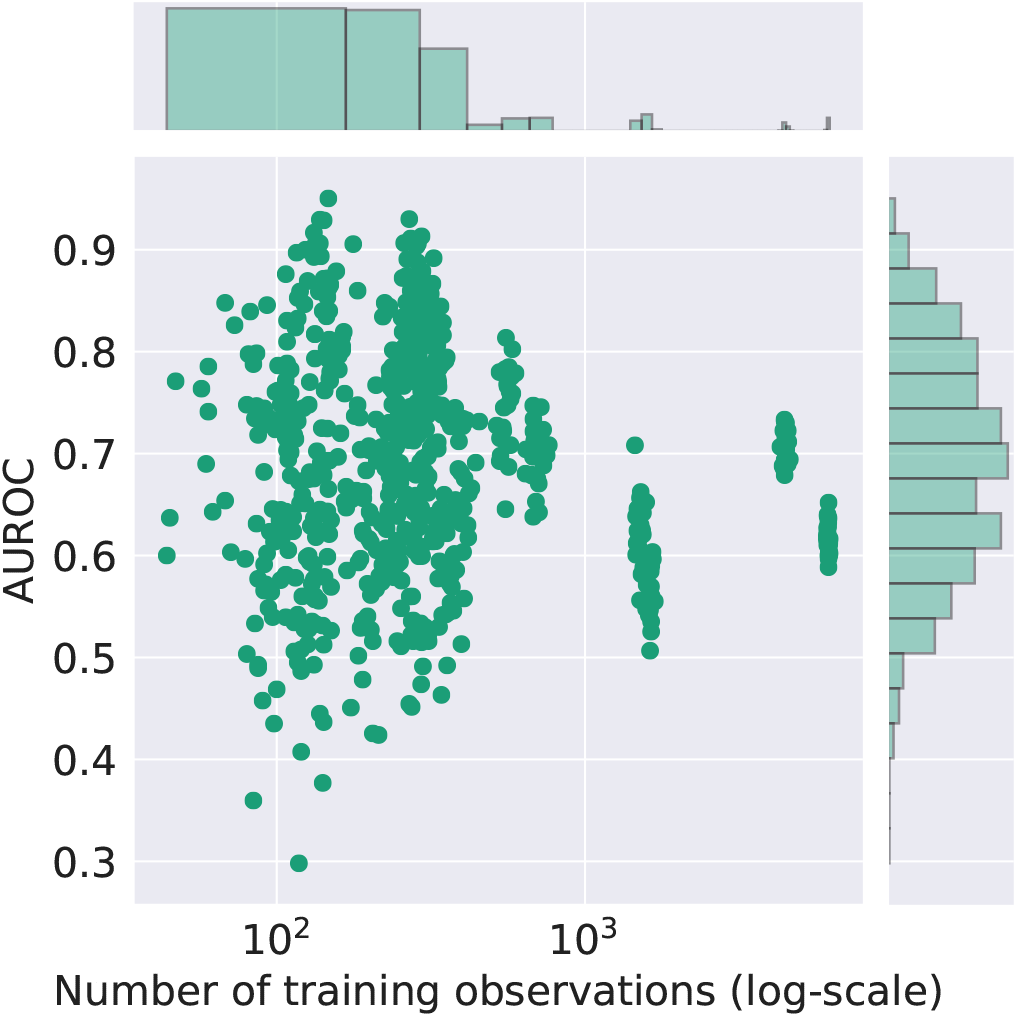
Scatter plot of per-epitope AUROC and training sample size for the ImRex CNN when trained on the beta chain dataset with shuffled negatives using 5 times repeated 5-fold CV.

**Fig. S10.**
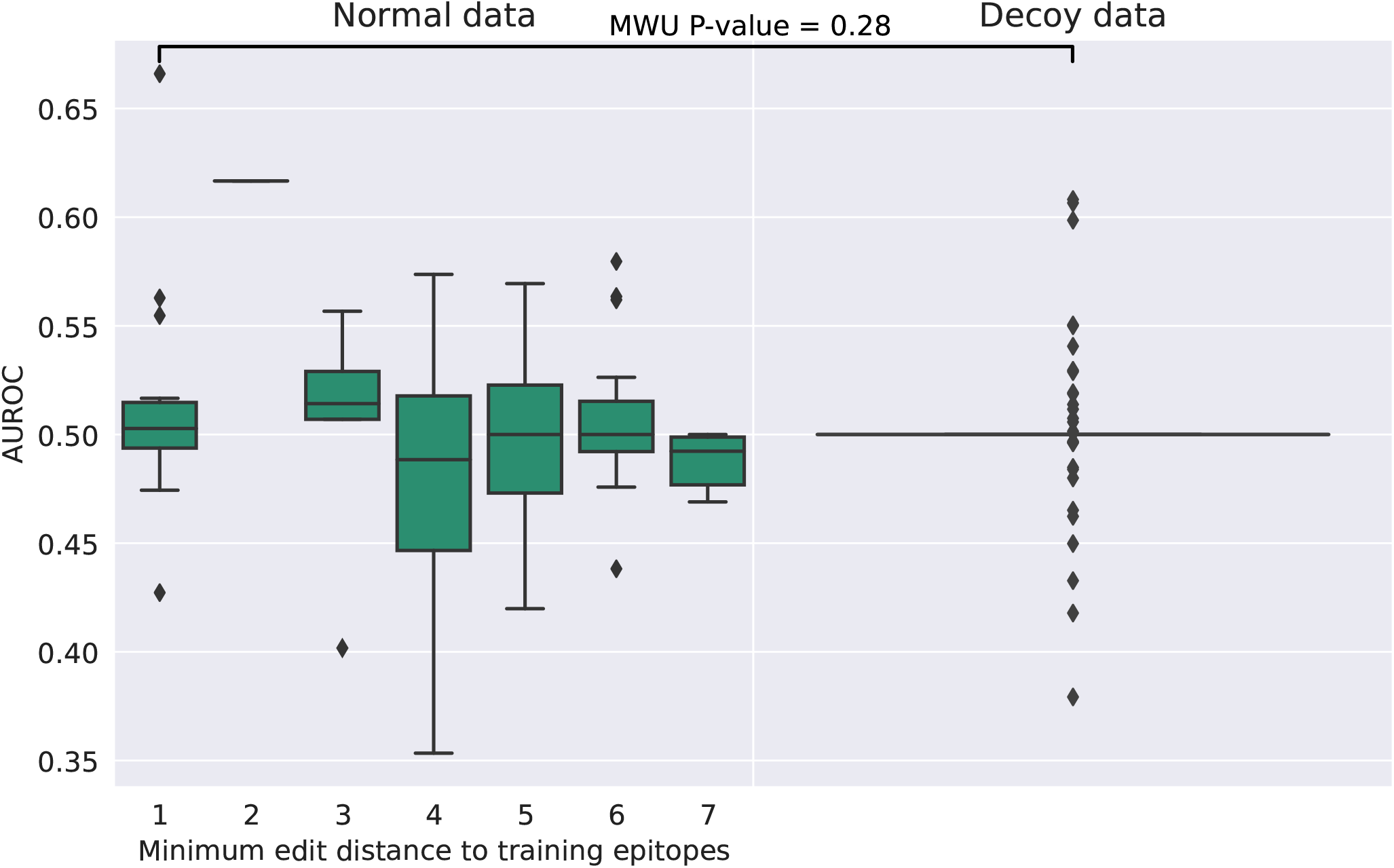
Box plots of per-epitope AUROC and minimum edit distance to epitopes in the training dataset for the dual input CNNs trained using 5 times repeated 5-fold CV on the downsampled beta chain dataset with shuffled negatives.

**Fig. S11.**
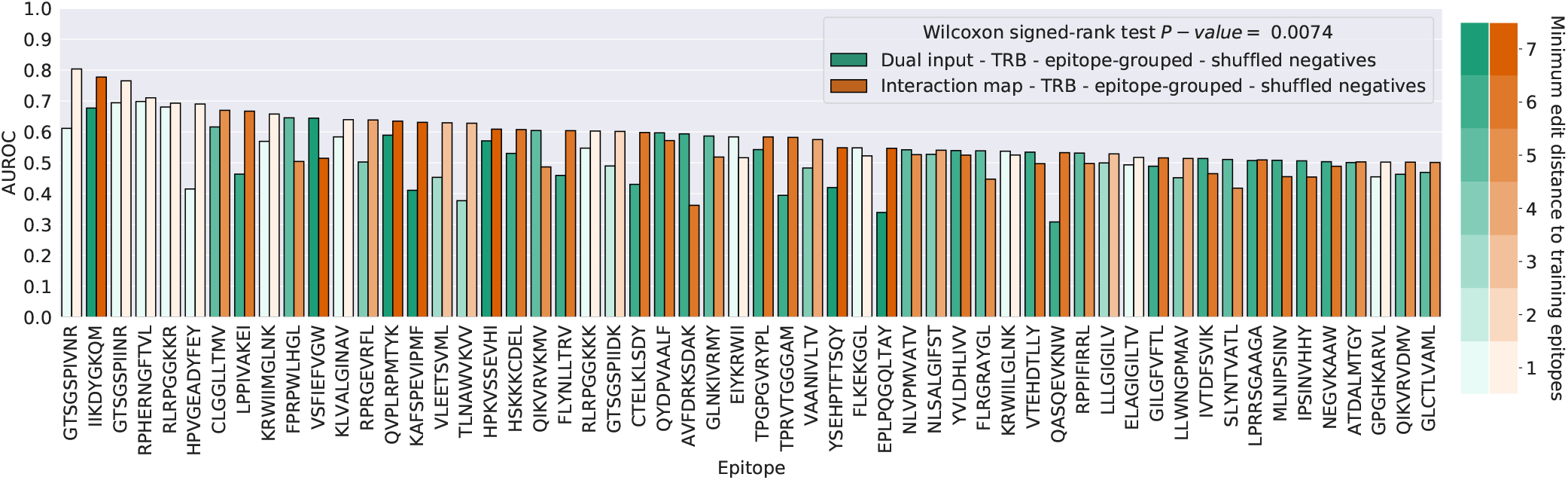
Ranking of the validation AUROC per epitope (≥ 30 examples and at least one model *AUROC* > 0.50) for the interaction map and dual input CNNs for a duplicate training run with 5-fold epitope-grouped CV using the downsampled beta chain dataset with shuffled negatives.

## Notes

https://github.com/pmoris/ImRex

https://doi.org/10.5281/zenodo.3973547

### Competing Interest Statement

The authors have declared no competing interest.

### Summary of Updates

New results including: improved methods of assessing per-epitope performance; use of decoy sequence data to establish baseline performances; investigation of relation between epitope distance and performance; comparison to single-epitope models (TCRex); validation on external dataset (McPAS-TCR); code and data repositories added.

